# The role of learned song in the evolution and speciation of Eastern and Spotted towhees

**DOI:** 10.1101/2023.05.15.540821

**Authors:** Ximena León Du’Mottuchi, Nicole Creanza

## Abstract

Oscine songbirds learn vocalizations that function in mate attraction and territory defense. Sexual selection pressures on these learned songs could accelerate speciation. The Eastern and Spotted towhees are sister species that diverged recently (0.28 Ma) but now have partially overlapping ranges with evidence of some hybridization; widespread community-science recordings of these species, including songs within their zone of overlap and from potential hybrids, enable us to investigate whether song differentiation might facilitate their reproductive isolation. Here, we quantify 16 song features to analyze geographic variation in Spotted and Eastern towhee songs and test for species-level differences. We then use random-forest models to measure how accurately their songs can be classified by species, both within and outside the zone of overlap. While no single song feature reliably distinguishes the two species, a random-forest model trained on 16 features accurately classified 89.5% of songs; interestingly, species classification was less accurate in the zone of overlap. Finally, our analysis of the limited publicly available genetic data from each species supports the hypothesis that they are reproductively isolated. Together, our results suggest that, in combination, small variations in song features may contribute to these sister species’ ability to recognize their species-specific songs.

## INTRODUCTION

In the Oscine songbirds, birdsong is a set of learned vocalizations used primarily for attracting mates and defending territories, and the characteristics of these songs vary widely across different species (Catchpole and Slater 1995). Song also varies within a species, with some differences in birdsong characteristics occurring at large spatial scales (Searfoss, Liu, and Creanza 2020). This long-range geographic variation could be influenced by a combination of genetic and environmental factors, such as adaptation to local habitats and ecological conditions (Podos and Warren 2007). Furthermore, under the assumption of isolation-by-distance due to spatially limited dispersal, songs are expected to accumulate differences gradually over time and become increasingly different with geographic distance (Rivera-Gutierrez et al. 2010). In vocal learners, such as songbirds, geographic variation can also accumulate through cultural transmission of learned vocalizations because of many factors, including, but not limited to, physical isolation and divergence in sexual selection (Podos and Warren 2007).

Divergence in traits that are under sexual selection is important for premating reproductive isolation as two species diverge (Price 2007), since mate preference based on these traits can limit interpopulation mating, which is beneficial if hybrid offspring have lower fitness (Ptacek 2000; MacDougall-Shackleton and MacDougall-Shackleton 2001; Slabbekoorn and Smith 2002; Lachlan and Servedio 2004; Podos 2007). For example, in a study of chorus frogs (*Pseudacris nigrita nigrita* and *Pseudacris triseriata feriarum*), the (unlearned) calls of the two species show significant differences in pulse rate and pulse number in sympatry, suggesting that these characteristics may have diverged via reproductive character displacement (Fouquette 1975). A similar pattern has been observed for a learned behavior: Ratcliffe and Grant (1985) found that although songs of *Geospiza fortis* and *Geospiza fulginosa* had very similar syllable structure and timing, individuals still preferred their own species’ songs in a playback experiment, suggesting that they are likely using subtle song differences to inform their mate choice. Since birds can use their songs for species recognition, these song differences can inhibit birds from choosing mates from different locations with less familiar song characteristics (Podos 2007; Price 2007). In theory, a rapid accumulation of learned song changes can lead to reproductive isolation, accelerating speciation (Lachlan and Servedio 2004).

Hybridization can occur when two genetically distinct populations come into contact with one another and mate (Barton & Hewitt 1985, 1989). Searching for a mate can be costly to females if conspecific males are difficult to find in a given area. Therefore, in a zone of overlap between two closely related species, females of the rarer species could have an increased tendency to hybridize with mates of the more common species, particularly if the signal of the heterospecific male is similar to that of the conspecific male (Price 2007). Stability of the hybrid zone can be maintained via spatial segregation of the two populations: as the hybrid zone is traversed, individuals of one species will become less common, leading them to mate more frequently with the other species and produce hybrids with reduced fitness (Price 2007).

In some hybrid zones, the hybrids between two species have high fitness and are not selected against, so the hybrid zone can be quite large (Grant and Grant 1992; Price 2007). However, cross-mating does not necessarily lead to the populations merging into one. For example, selection that favors the prevalent phenotype of a given area constrains the width of the hybrid zone because individuals that disperse in the range of the other species could exhibit inferior fitness (Moore and Price 1993; Price 2007). Further, separation of species can be maintained by reinforcement if the strength of selection is greater for the characteristics that fall in the extremes rather than those of the hybrids (Price 2007), such that premating isolation is strengthened. In other words, if sexual selection favors more extreme characteristics, then hybrids will have lower mating success. For example, a study of the same two species of chorus frogs (*Pseudacris nigrita* and *Pseudacris feriarum*) showed evidence that hybrid fitness was significantly reduced, suggesting that reinforcement, via sexual selection against hybrid males, is the primary mechanism behind the reproductive character displacement in these species (Lemmon and Lemmon 2010).

The Spotted towhee (*Pipilo maculatus*) and the Eastern towhee (*Pipilo erythrophthalmus*) are sister species of the Passerellidae family of oscine songbirds. It is estimated that the Eastern and Spotted towhees diverged 280,000 years ago (Hirase et al. 2016; DaCosta et al. 2009). Evidence from the geological history of the Great Plains, in combination with patterns of the current geographic distribution of the towhees and their character gradients, suggest that present hybridization of these sister species in the Great Plains is a result of secondary contact that occurred after the Pleistocene glaciation fragmented their breeding range and accelerated speciation (Sibley and West 1959; Hirase et al. 2016; Price 2007). They were previously considered one species, the Rufous-sided towhee, and the idea that the Spotted and Eastern towhees should be considered separate species has been contentious for many years. These sister species were separated into their respective groups primarily due to distinctions in their geographic distribution, genetic data, and morphological characteristics such as plumage, sexual dichromatism, and eye color (Ball and Avise 1992). Secondarily, early studies noted some differences between songs in the eastern and western populations (Sibley and West 1959).

The breeding range of the Spotted towhee lies on the west side of the United States extending into Mexico and southern Canada, while the breeding range of the Eastern towhee lies on the east side of the United States and southern Canada. Hybridization of the Spotted and Eastern towhees has been reported in the Great Plains area, particularly in Nebraska, where towhees had been noted as exhibiting ‘intermediate’ songs (Sibley and West 1959). Further, Sibley and West (1959) used plumage as an index for hybridization by scoring birds in the Great Plains based on spotting of the wing coverts (males) and color of the head and back (females). Although hybridization occurs, they are still able to maintain themselves as separate species. While these species share many similarities in their vocalizations, subtle but consistent differences in their songs could allow individuals to distinguish between the different species and identify potential conspecific mates, allowing separation of species to be maintained, a form of behavioral premating isolation.

The typical song of both the Spotted towhee and Eastern towhee is a loud and clear “drink-your-tea”, composed of short introductory notes and a fast trill (**Fig. 1)**. Previous studies suggest that the Eastern towhees tend to have songs that are more variable and complex, with a greater number of syllables, while the Spotted towhees tend to have fewer syllables or no introductory syllables with a faster trill (Borror 1959; Kroodsma 1971). In this study, we aim to distinguish any geographic patterns in the song characteristics of the Spotted towhee and Eastern towhee, which have not been quantified between the two species. By analyzing the songs of each species individually as well as together as a two-population cline, we aim to understand if there are song patterns that are continuous across the geographic ranges of the Eastern and Spotted towhees or if there are discontinuities in the geographic distribution that clearly distinguish the songs of the two species from one another. Additionally, we use machine learning to investigate whether the two species differ in their song characteristics such that birds could potentially use multiple song features to distinguish between conspecific and heterospecific individuals.

**Figure 1.**
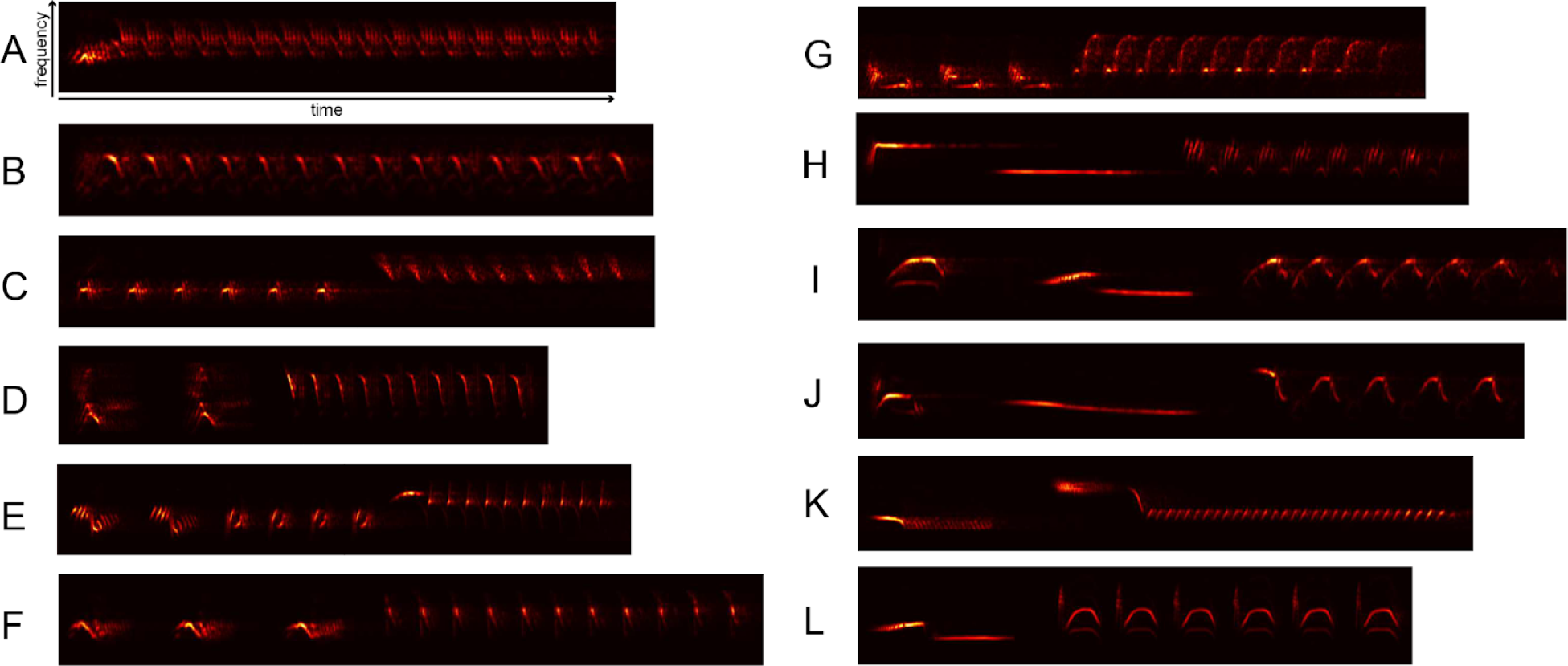
Example spectrograms of (**A-F**) Spotted towhee songs and (**G-L**) Eastern towhee songs. The repeated syllable at the end of each song is often called the ‘trill’.

## METHODS

### Generating a database of song bouts

We downloaded recordings of Spotted and Eastern towhees from the Macaulay Library (Cornell Lab of Ornithology 2009) by requesting access to the relevant song files, as well as from Xeno-canto (Planque and Vellinga 2005) using the WarbleR package in R to query the database (Araya-Salas and Smith-Vidaurre 2017). We filtered the recordings to eliminate those without song by specifying “Sounds/Type = Song”. We also requested recordings marked as “Spotted x Eastern Towhee (hybrid)” and “Spotted/Eastern Towhee (Rufous-sided Towhee)” from the Macaulay Library to analyze separately as a “hybrid/unsure” category. For further analysis, we constructed a spreadsheet of metadata for each recording, which included species, date, location, latitude, longitude, and recordist. To minimize the chance of including repeated song recordings of the same bird, we discarded any duplicate entries with the same recordist, date, and location. For recordings that named a location but not geographic coordinates, we estimated the latitude and longitude by identifying the location on a map. Each recording file was opened in Audacity version 3.1.3 (https://www.audacityteam.org/) to manually extract one or more bouts depending on the number of unique song types from each recording. Files from Xeno-canto are generally stored in MP3 format and files from Macaulay Library in WAV format; we exported each bout as a WAV file with a sampling rate of 44100 Hz to standardize the recordings for analysis. We considered the breeding season to be between April and August based on the “Breeding” section of the Birds of the World database (Bartos and Greenlaw 2020; Greenlaw 2020), and we conducted analyses both on all recordings (main text) and on breeding season recordings (**Table S1**, **Supplementary Materials**).

To assign the zone of overlap between the ranges of the two species, for each degree longitude we plotted the number of recordings of each species over the total number of recordings of both species (restricted to the breeding season). At the most eastern and western longitudes, the recordings were all from Eastern and Spotted towhees, respectively. Based on our findings, we chose to focus on the area between 102°W and 91°W; which contained the majority of the species’ range overlap (**Fig. S1**). In addition, we downloaded from eBird all sightings of putative hybrids (eBird query: “Spotted x Eastern Towhee (hybrid) - Pipilo maculatus x erythrophthalmus”) during the breeding season and plotted their locations; we observed that reported hybridization events primarily occurred in this region (**Fig. S2**). We also determined the approximate frequency of hybridization in that area during recent years by limiting the sightings to those in breeding seasons between 2013 and 2023 and calculating the ratio of hybrid sightings over the number of Spotted towhee or Eastern towhee sightings, filtering out those from the same date, latitude, and longitude.

### Song analysis (syllable features and syllable segmentation)

To begin analyzing the features of the syllables within these bouts, we used the Chipper software (Searfoss, Pino, and Creanza 2020), which was designed to facilitate syllable segmentation and analysis of field recordings with different levels of background noise. This software parses each song into syllables by identifying periods of sound separated by silences, and then allows the user to enable high-pass and low-pass filters and to modify the segmentation when syllables have been incorrectly parsed, such as when background noise occurs during the inter-syllable silences (**Fig. 2**). To prevent lower-amplitude syllables from being segmented incorrectly, we normalized the amplitude across each song. Additionally, we adjusted the signal-to-noise threshold, minimum syllable duration, and minimum silence duration to most accurately define the syllables in the song. If any syllables appear to be parsed incorrectly after this procedure, Chipper allows the user to manually modify the syllable segmentation. If, when visualized in Chipper, we determined that the song overlapped with other birds singing or had excessive background noise, we removed the recording from the analysis. We provide the analyzed song bouts and analysis parameters used in Chipper to extract song features, our code for statistical analysis and visualization of the song feature data, and metadata about each song recording at https://github.com/CreanzaLab/TowheeAnalysis.

**Figure 2.**
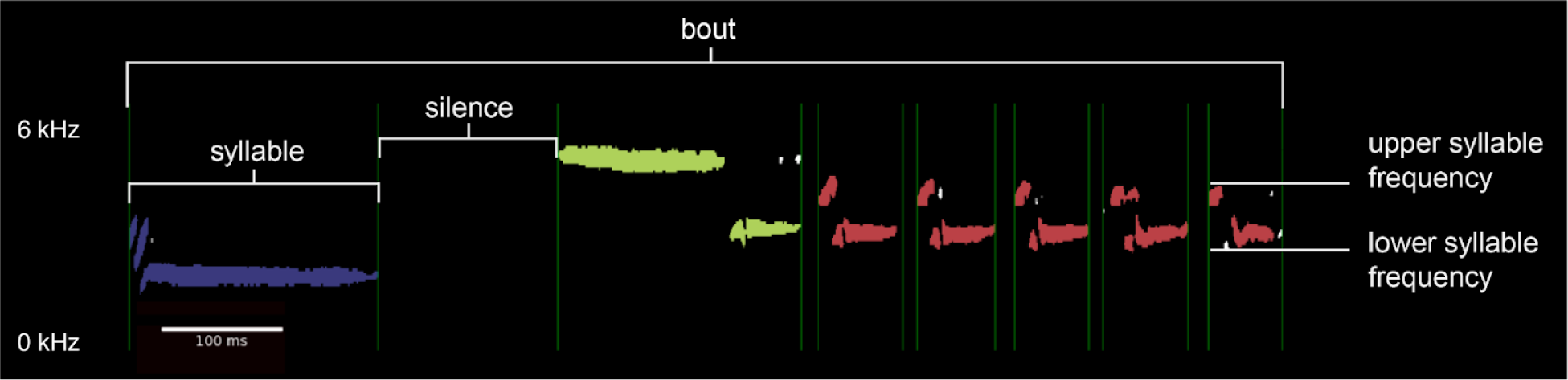
Spectrogram of Eastern towhee song illustrating how we categorized syllable, silence, bout, maximum syllable frequency, and minimum syllable frequency. The color of the syllable indicates its unique syllable type. Spectrogram generated from Macaulay Library recording 52506381, https://macaulaylibrary.org/asset/52506381.

After segmentation, we examined song bouts in Chipper to estimate a noise threshold, or the minimum number of matrix elements in the spectrogram that a note must contain in order to not be discarded as noise (Spotted towhee = 87, Eastern towhee = 70, hybrid/unsure = 65), and syllable similarity threshold, or the percent syllable overlap that determines whether two syllables are considered the same or different (Spotted towhee = 26.6, Eastern towhee = 25.5, hybrid/unsure = 25.0). We then ran the song analysis function in Chipper, which uses the spectrogram and the syllable segmentation data to provide automated measurements of numerous song features, which were used for statistical analysis. We used 16 song-feature outputs from Chipper: average syllable lower frequency (Hz), number of unique syllables, bout duration (ms), smallest syllable duration (ms), number of syllables per bout duration (1/ms), maximum syllable frequency (Hz), average syllable frequency range (Hz), minimum syllable frequency (Hz), largest syllable duration (ms), number of syllables per number of unique syllables, average syllable duration (ms), smallest syllable frequency range (Hz), number of syllables, overall syllable frequency range (Hz), largest syllable frequency range (Hz), and average syllable upper frequency (Hz). These features were analyzed across 1427 Spotted towhee song bouts, 2088 Eastern towhee song bouts, and 31 “hybrid/unsure” song bouts.

### Statistical analysis of song features

To investigate the geographic variation in birdsong of the Spotted and Eastern towhees, we first determined that the song features were not normally distributed, so we used the natural log of each feature. For this and subsequent analyses, we eliminated songs that had “NA” for one or more song features after the log transformation, which occurred when Spotted towhee recordings were parsed as one syllable because the trill could not be resolved into separate syllables. Our analysis thus contained a total of 1427 Spotted towhee samples and 2088 Eastern towhee samples and data for 16 song features. To visualize the data, we plotted each log-transformed feature against the longitude and latitude coordinates of its recording.

To test whether a directional pattern exists between the longitude and latitude and the song features of the Spotted towhees and the Eastern towhees, we used Spearman rank correlation to measure the strength and direction of association across the 3515 Spotted and Eastern towhee song bouts. We used the Bonferroni multiple hypothesis correction to adjust our alpha threshold for statistical significance for 16 song features and analysis of longitude and latitude (ɑ_adjusted_=0.00156). Additionally, we used a Wilcoxon rank-sum test to test for species-level differences between Eastern and Spotted towhees in song features using Bonferroni multiple hypothesis correction for 16 song features (ɑ_adjusted_=0.003125).

### Principal components and Procrustes analyses of song feature data

We used a Principal Component Analysis (PCA) to reduce the dimensionality of the dataset and visualize the variation in the song data, and we measured the proportion of variance explained by each principal component (PC). Using the ‘prcomp’ function in the stats package in R, we centered and scaled the data and performed the PCA. We assessed whether the two species could be distinguished in PC space by testing a simple linear classifier using the ‘lda’ function in the MASS package in R. This type of Discriminant Function Analysis finds a linear function that best discriminates the ‘Species’ classification of the song recordings based on PC1 and PC2. In other words, this classifier finds a line through the two-dimensional PC-space such that songs on one side of the line are more likely to be from Spotted towhees and on the other side of the line are more likely to be from Eastern towhees, indicating the separability of the species after PCA (Armstrong *et al*. 2021). Further, to determine whether the PCA showed geographic structuring, we followed up with a Procrustes analysis using PC1 and PC2 from the PCA of our song data. For this analysis, the matrix of PC scores was rotated and transformed onto the target matrix containing the longitude-latitude coordinates of the samples, such that the sum of squared distances between the corresponding points of the transformed and target matrix was minimized and plotted on a map using the ‘procrustes’ function from the vegan package (Oksanen et al. 2023) in R.

### UMAP visualization

We used a Uniform Manifold Approximation and Projection (UMAP) as another method of reducing dimensionality and visualizing variation in our song-feature data. We rescaled the data using the same technique as in the PCA, and we used the ‘umap’ function in the umap package in R to generate a two-dimensional projection of the data. Since most of the points occupy a single cluster in the resulting UMAP projection, we then tested a simple linear classifier as above to determine how well the songs could be separated by species in the UMAP projection. We included the sample of 31 “hybrid/unsure” towhees in the UMAP analysis, but we excluded these points when we tested the species classifier.

### Machine learning classifier

We used a random forest model (RFM) to assess how accurately Spotted and Eastern towhees can be distinguished by their song features, using the ranger package (Wright and Ziegler 2017) in R. The model was trained on the 16 song features from the training set and multiple decision trees were averaged to make predictions of the species identity in the test set (**Fig. 6**). We used our random forest algorithm to train two models: 1) a model trained on a geographically unbiased subset of the entire data set and 2) a model trained on a subset of song bouts obtained from the non-overlap zone. For the former model, we obtained a random subsample (75% for training and 25% for testing) of the entire data set, and we then downsampled the training set to obtain a balanced subsample of Eastern towhees equal to the number of Spotted towhee song bouts (N_Spotted_towhee_=1062 and N_Eastern_towhee_=1062). We tested the model on 879 song bouts, none of which were included in the training set. For the latter model, we separated the song bouts recorded in the zone of overlap from those in the non-overlap regions, and trained only on a subset of the non-overlap song bouts, retaining a sample size equal to that of the zone of overlap for testing (**Fig. 6A**). Once again, our training set was downsampled, such that we obtained a balanced subsample of Eastern towhees equal to that of Spotted towhees in our training set (N_Spotted_owhee_=1062 and N_Eastern_towhee_=1062). We tested this model on both a sample of towhees from the non-overlap zones (N=299), an equally sized sample of towhees from the zone of overlap (N=299), and the sample of “hybrid/unsure” towhees (N=31; **Fig. 6B and C; Fig. S5**). For both models, we created a 10-fold cross-validation control and tuned the hyperparameters using a 10 x 16 grid using the caret package (Kuhn 2008) in R, which allows us to evaluate every possible combination of the number of predictors to randomly sample at each split and the minimum number of samples needed to keep splitting nodes using Gini impurity to split nodes. Additionally, we tested the models using both 100 and 1000 trees and obtained similar results (**Table S2**). Our code for all analyses is available at https://github.com/CreanzaLab/TowheeAnalysis.

### Genetic analysis

We downloaded fasta files of sequences of the cytochrome oxidase subunit I regions of the Spotted and Eastern towhee mitochondrial genomes available in the Barcode of Life Data Systems (BOLD) database and the NCBI database. We obtained a total of 18 Spotted towhee and 5 Eastern towhee sequences. We aligned these sequences using MAFFT (Katoh, Asimenos, and Toh 2009), and we used a pairwise F_ST_ (fixation index) in Arlequin (Excoffier and Lischer 2010) to quantify the level of genetic differentiation between the two species by measuring the proportion of genetic variation that is due to differences between populations. It also measures the extent to which individuals within populations are similar to one another, where a larger F_ST_ describes a greater difference in allele frequencies within a population. We also did an Analysis of Molecular Variance (AMOVA) to quantify the proportion of genetic variation that is due to differences between versus within populations.

### PCA and Procrustes analysis of genetic data

Using the glPCA function from the adegenet package (Jombart 2008) in R, we ran a PCA on the aligned fasta files from our genetic analysis. We followed with a Procrustes analysis using PC1 and PC2, and we transformed the two-dimensional plot of the first two principal components onto the target geographic plot of the longitude-latitude coordinates of the samples using the ‘procrustes’ function from the vegan package (Oksanen et al. 2023) in R. We then performed a resampling test using the ‘protest’ function in the vegan package.

## RESULTS

After gathering all song recordings of Eastern and Spotted towhees from the largest public repositories of natural sounds, Xeno-canto and Macaulay Library, and eliminating recordings that lacked song or had excessive background noise, we were able to analyze 3515 total song bouts: 2088 song bouts from 1563 individual Eastern towhees and 1427 song bouts from 874 individual Spotted towhees, in addition to 31 total song bouts from 27 recordings from the “hybrid/unsure” category. For each song bout, we segmented the song into syllables, defined as periods of sound separated by periods of silence, and we used the song-analysis software Chipper (Searfoss, Pino, and Creanza 2020) to automatically extract 16 syllable and song features. In the Spotted towhee, several samples had very rapid trills that could not be further parsed into syllables (i.e. there were no periods of silence within the trill (**Fig. 1**)) and, thus, were considered to be a single syllable. Therefore, a subset of the Spotted towhee samples with one syllable in the bout were recorded from birds with a rapid trill that could not be further subdivided by silences.

To estimate the frequency of hybridization events in the zone of range overlap (between −91° and −102° longitude), we estimated the number of unique sightings of hybrid towhees during the breeding season divided by the number of unique Eastern and Spotted towhee sightings. We estimated unique sightings of 314 unique hybrids in the same region as sightings of 71,521 Eastern towhees (0.44%) and 20,197 Spotted towhees (1.6%).

### Statistical analysis of song features

Statistical analysis revealed several patterns in the song features of the Spotted and Eastern towhees. Spearman rank correlation showed statistical significance (*p* < 0.00156) in 14 syllable features across longitude and 11 features across latitude (**Table 1**). Stronger correlations (ρ > 0.400) existed in song features across the longitude gradient (e.g. bout duration, number of unique syllables, average syllable upper frequency, and smallest syllable frequency range, **Table 1**, **Fig. 3A-E**). A Wilcoxon rank-sum test showed statistical significance (*p* < 0.003125) in all but 2 of the song features (**Fig. 3F-J**). We observed qualitatively similar results when we restricted the analysis to only songs that were recorded during the breeding season: the same set of song features showed significant associations with latitude and longitude (**Table S1**).

**Figure 3.**
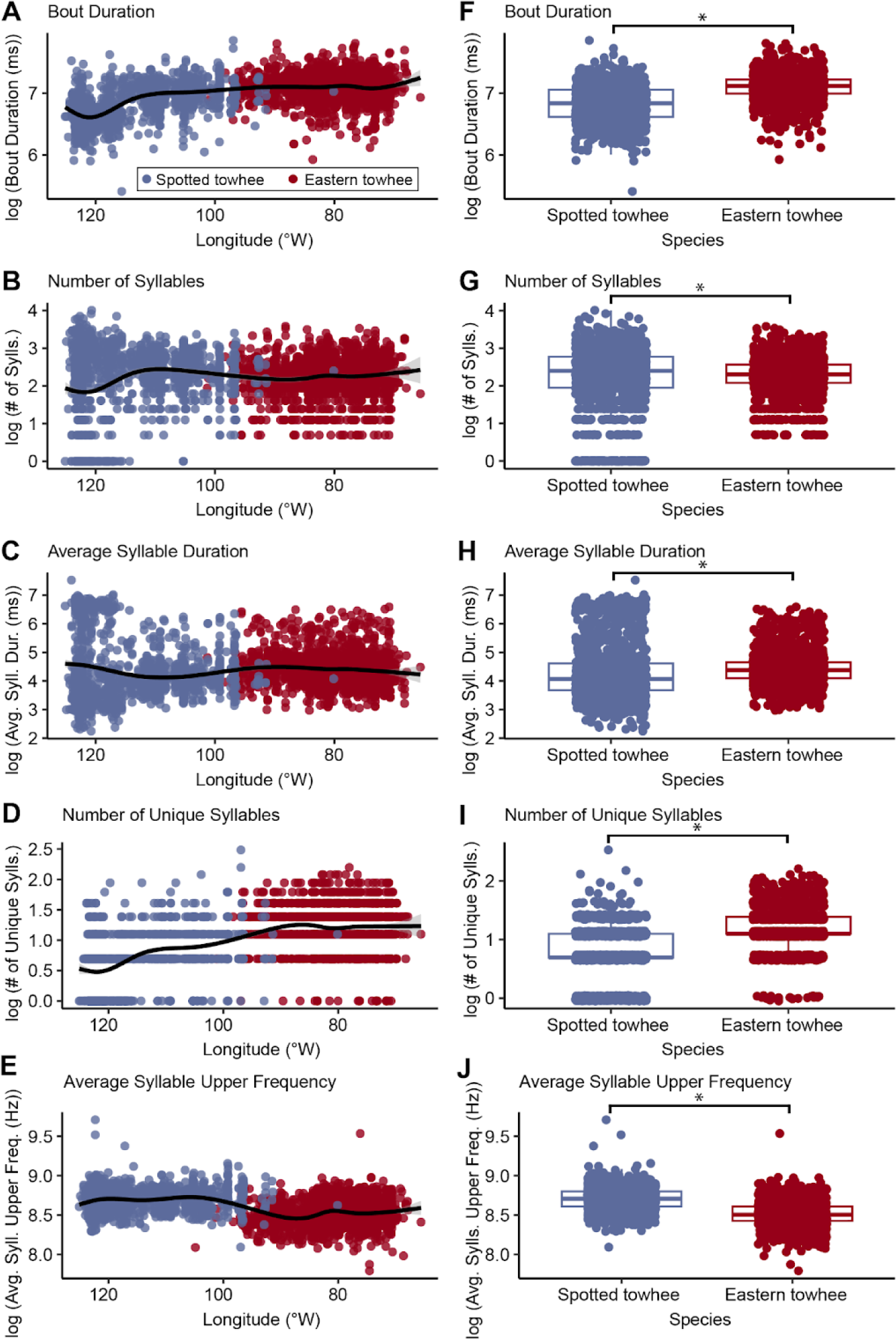
Spearman rank correlation (**A-E**) and Wilcoxon’s rank sum test (**F-J**) of five song features in songs of Spotted towhees and Eastern towhees (N_total_bouts_=3515; N_Spotted_towhee_=1427; N_Eastern_towhee_=2088). (Spearman: ɑ_adjusted_=0.00156; Wilcoxon: ɑ_adjusted_=0.003125). The black line on panels A-E represents a generalized additive model-fitting method. Asterisks (*) in panels **F-J** indicate statistical significance; Spearman’s rank correlations were significant in panels **A**, **C**, **D**, and **E**; the number of syllables (panel **B**) was not significantly associated with longitude.

**Table 1.** Spearman rank correlation (ɑ_adjusted_=0.00156) and Wilcoxon rank-sum test (ɑ_adjusted_=0.003125) of Spotted towhee and Eastern towhee song features (N=3515; N_Spotted_towhee_=1427; N_Eastern_towhee_=2088).

| Song Feature | Spearman |  |  |  |  |  | Wilcoxon |  |
| --- | --- | --- | --- | --- | --- | --- | --- | --- |
|  | Latitude |  |  | Longitude |  |  |  |  |
| | S | p-value | Spearman's $\rho$ | S | p-value | Spearman's $\rho$ | W | p-value |
| bout duration (ms) | 7397584966 | 0.192 | -0.022 | 3748450720 | 4.40E-204 | 0.482 | 654894 | 1.18E-175 |
| number of syllables | 7827213145 | 1.35E-06 | -0.081 | 7359761359 | 0.319 | -0.017 | 1626277 | 3.67E-06 |
| number of syllables per bout duration (1/ms) | 7622436939 | 1.64E-03 | -0.053 | 8589473544 | 6.15E-29 | -0.187 | 1924310 | 5.88E-49 |
| largest syllable duration (ms) | 6771885315 | 1.33E-04 | 0.064 | 5522445809 | 4.39E-46 | 0.237 | 896183 | 8.97E-90 |
| smallest syllable duration (ms) | 6731656553 | 3.30E-05 | 0.070 | 7423283837 | 0.129 | -0.026 | 1474316 | 0.601 |
| average syllable duration (ms) | 6680317932 | 4.79E-06 | 0.077 | 6266818563 | 1.36E-15 | 0.134 | 1135866 | 4.61E-33 |
| number of unique syllables | 6992434693 | 0.044 | 0.034 | 3453642343 | 6.34E-246 | 0.523 | 534212.5 | 4.72E-244 |
| number of syllables per number of unique syllable | 7832890160 | 1.07E-06 | -0.082 | 9075790484 | 7.82E-53 | -0.254 | 2089206 | 1.10E-91 |
| average syllable upper frequency (Hz) | 8248493705 | 9.26E-17 | -0.140 | 10162112044 | 4.08E-138 | -0.404 | 2438590 | 3.02E-226 |
| average syllable lower frequency (Hz) | 7277970268 | 0.744 | -0.006 | 8380047396 | 4.99E-21 | -0.158 | 1864955 | 6.12E-37 |
| maximum syllable frequency (Hz) | 8436848622 | 4.89E-23 | -0.166 | 7929152107 | 1.42E-08 | -0.095 | 1880803 | 5.53E-40 |
| minimum syllable frequency (Hz) | 7183062081 | 0.652 | 0.008 | 10128201861 | 1.09E-134 | -0.399 | 2204676 | 2.00E-129 |
| overall syllable frequency range (Hz) | 8398422023 | 1.15E-21 | -0.160 | 6640075404 | 9.34E-07 | 0.083 | 1548046 | 0.049 |
| largest syllable frequency range (Hz) | 8230917361 | 3.14E-16 | -0.137 | 8329654663 | 2.49E-19 | -0.151 | 1923190 | 1.01E-48 |
| smallest syllable frequency range (Hz) | 7901898144 | 5.13E-08 | -0.092 | 10290529942 | 1.32E-151 | -0.422 | 2393045 | 2.25E-205 |
| average syllable frequency range (Hz) | 8075151267 | 6.12E-12 | -0.116 | 9367136726 | 4.24E-71 | -0.294 | 2187303 | 3.26E-123 |

### PCA and Procrustes analysis of song data

A PCA of the song data revealed considerable overlap in the song variation captured by PC1 and PC2. Individual samples in both the range of overlap and non-overlap were scattered throughout PC space; if the individuals in the zone of overlap had songs that showed greater species-level differentiation, we would expect that these points would have greater separation in PC space than those in the zone of non-overlap. PC1 explained 35.4% of the variance, with the highest loading being average syllable duration. PC2 explained 24.7% of the variance, with average syllable upper frequency most closely associated to it. With a Linear Discriminant Analysis, we could predict the species of a song in two-dimensional PC-space with 73.6% accuracy. A Procrustes analysis comparing the first two principal components of the song data to the latitude-longitude coordinates of the recording location indicated that, despite the extensive overlap between the two species, there is significant geographic signal in the song data (Procrustes rotation 0.26, *p* < 0.001, **Fig. 4A-C**).

**Figure 4.**
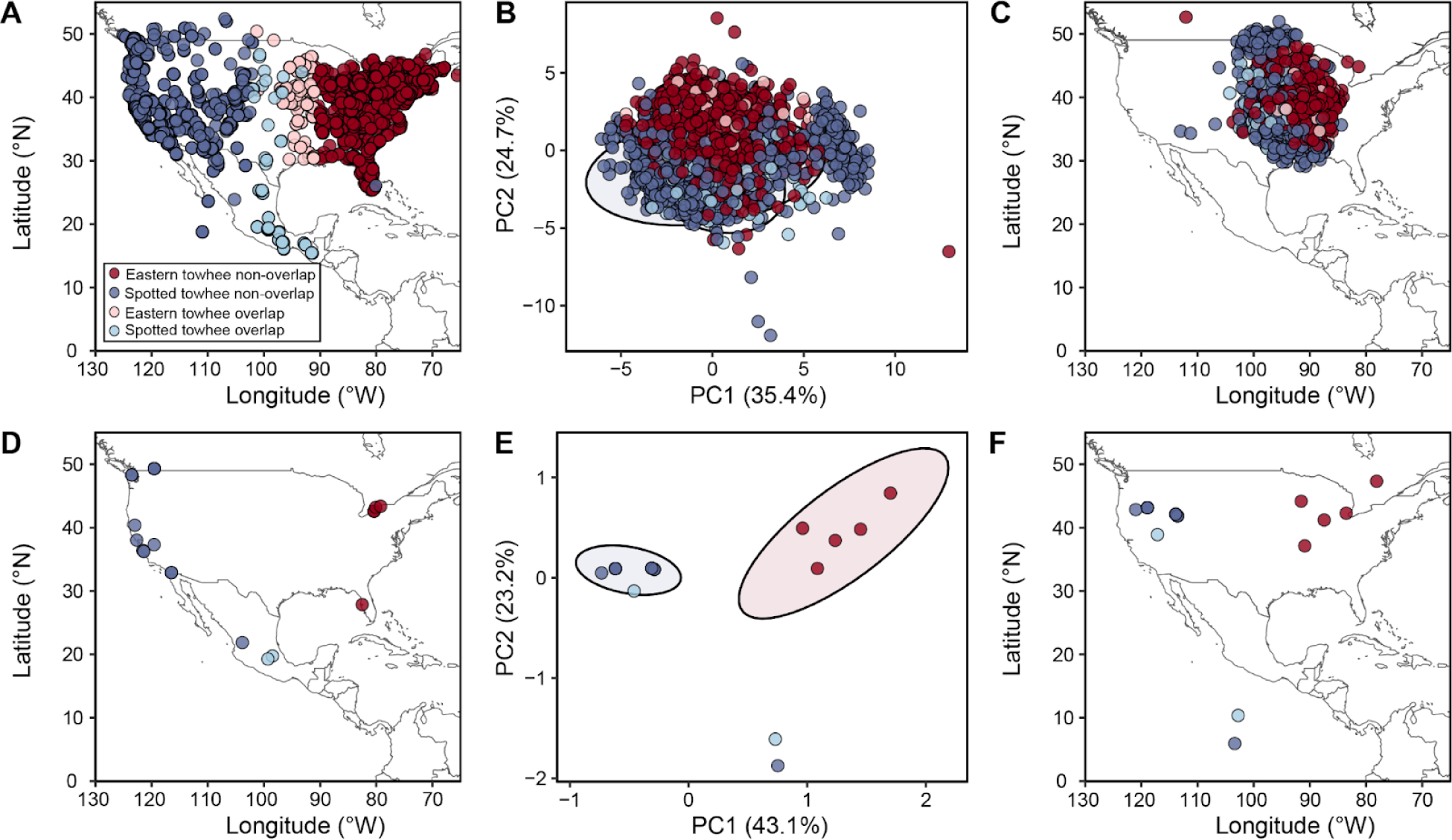
Spatial distribution of Spotted towhee and Eastern towhee song (**A-C**) and genetic (**D-F**) data. (**A**) Distribution of song recordings in North America (N_total_recordings_=2788; N_Spotted_towhee_=1069; N_Eastern_towhee_=1719). (**B**) Principal component analysis (PCA) of song bouts using 16 song features (N_total_bouts_=3515; N_Spotted_towhee_=1427; N_Eastern_towhee_=2088). (**C**) Procrustes analysis of song data using PC1 and PC2 from the PCA in panel B (N_total_bouts_=3515; N_Spotted_towhee_=1427; N_Eastern_towhee_=2088). (**D**) Distribution of genetic sequences obtained from the Barcode of Life Data Systems database and NCBI (N_total_=23 ; N_Spotted_towhee_=18; N_Eastern_towhee_=5). (**E**) PCA using single nucleotide polymorphisms of aligned sequences of the cytochrome oxidase subunit I regions of the mitochondrial genome (N_total_=23 ; N_Spotted_towhee_=18; N_Eastern_towhee_=5). (**F**) Procrustes analysis of genetic data using PC1 and PC2 from the PCA analysis in panel E (N_total_=23 ; N_Spotted_towhee_=18; N_Eastern_towhee_=5). Ellipses indicate 95% confidence intervals.

### UMAP visualization

Projecting the song-feature data with a UMAP analysis showed that the songs of the Eastern and Spotted towhees did not form two species-specific clusters, but instead made one large cluster with Eastern towhee songs overrepresented on one side and Spotted towhee songs on the other side (**Fig. 5**); two much smaller clusters contained predominantly Spotted towhee songs and a third contained a mixture of both species. As in the PCA (**Fig. 4**), songs sampled from the range of overlap were scattered throughout the UMAP projection. The species classification of each song could be better discriminated in the UMAP projection than the PCA plot; with a simple linear partitioning of the two-dimensional UMAP projection with n_neighbors=15 and min_dist=0.1, we could predict the species of a song with 84.4% accuracy. We tested values of n_neighbors up to 50 and values of min_dist up to 0.9 and found very similar prediction accuracies (80.7%–84.5%). The “hybrid/unsure” recordings were located primarily towards the edges of the UMAP projection (**Fig. 5**). The same analysis using data from samples recorded only during the breeding season showed similar results (**Fig. S3**).

**Figure 5.**
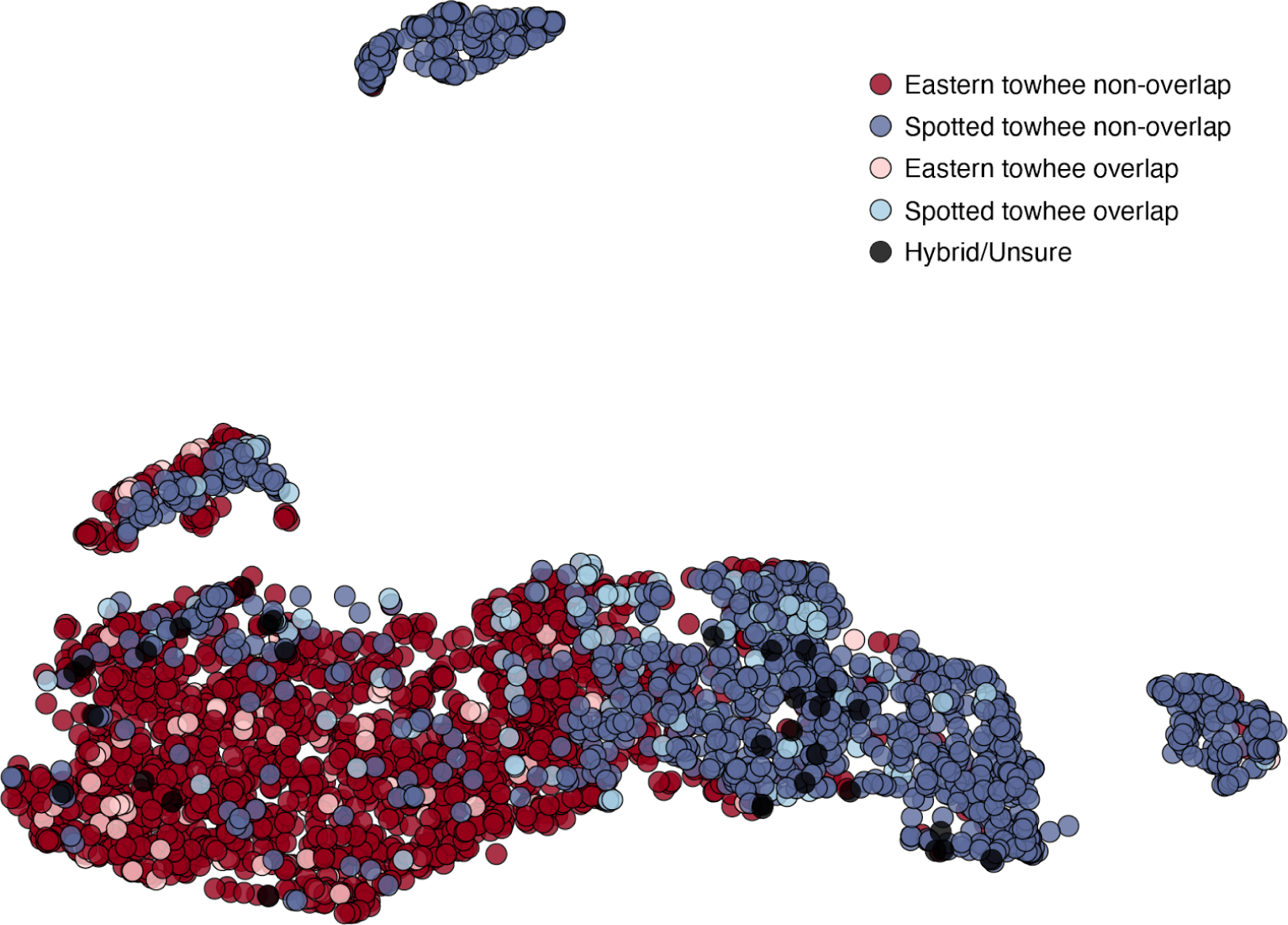
UMAP projection of Eastern and Spotted towhee song-feature data. Each point represents an analyzed song bout (N_total_bouts_=3515; N_Spotted_towhee_=1427; N_Eastern_towhee_=2088), with Eastern towhee songs shown in shades of red and Spotted towhee songs in shades of blue. The lighter colors represent recordings from the zone of species overlap. Black dots indicate the 31 recordings from individuals that were classified as potential hybrids (“hybrid/unsure”).

### Machine learning classifier

Our random forest model trained on samples from the entire geographic range had an accuracy of 89.5% when tested on a subset of song samples that had been withheld from training. The most important feature in the decision tree was the average syllable upper frequency. Our model trained on the samples from the zone of only non-overlap was tested on a withheld subset of samples from the zone of non-overlap (299 song bouts) and on all samples from the zone of overlap (299 song bouts) which had an accuracy of 89.3% and 82.6%, respectively. The most important feature was the largest syllable duration. When we used the model trained on only the samples from the zone of non-overlap to predict the species classification of the samples from “hybrid/unsure” towhee recordings, it predicted 17 Spotted towhees and 14 Eastern towhees (**Fig. 6C; Fig. S5**)

**Figure 6.**
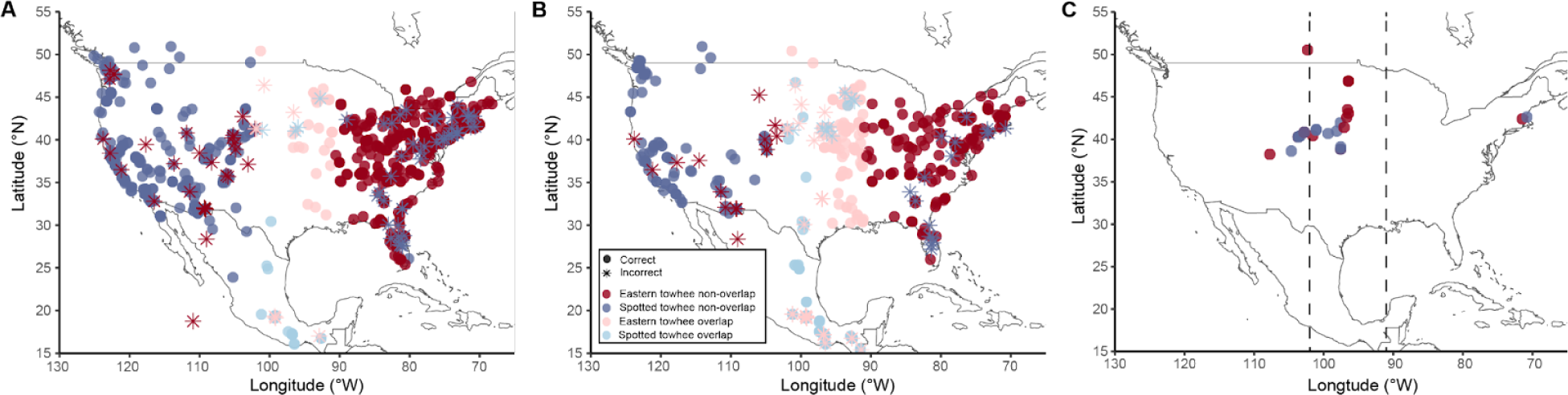
Geographic distribution of random forest model predictions of species identity based on a model trained on 16 song features from samples of Spotted towhees and Eastern towhees. (**A**) We trained a model on song data from the entire geographic range of both species (N_Spotted_towhee_=1062; N_Eastern_towhee_=1062) and tested how well it predicted the species identification of a subset of all song samples (N_test_=879; accuracy = 89.5%). (**B**) We then trained a second model on a subset of samples obtained only from the non-overlap zone (N_Spotted_towhee_=1062; N_Eastern_towhee_=1062) and tested it on a random subsample of song bouts from both the zone of non-overlap (N_test_nonoverlap_=299; accuracy = 89.3%) and the zone of overlap (i.e. 102°W - 91°W; N_test_overlap_=299; accuracy = 82.6%). (**C**) We used the same model from panel **B** to predict species identity of song bouts from recordings of “hybrid/unsure” towhees (N_predict_=31). The model predicted that 17 of these “hybrid/unsure” recordings were Spotted towhees and 14 were Eastern towhees, with no discernable longitudinal gradient in the predictions. The dotted line represents the zone of overlap determined by the co-occurence of Eastern towhee and Spotted towhee song recordings (102°W - 91°W).

### PCA and Procrustes analysis of genetics data

A PCA using mtDNA sequences revealed 3 different clusters. Eastern towhees formed their own cluster, while the Spotted towhees were split into two clusters: one group that contained two individuals from Mexico and a second group that included all individuals from the United States and Canada, along with one individual from Mexico. Eastern and Spotted towhees cluster separately, suggesting that they are genetically different, at least using SNPs from COI mtDNA sequences. PC1 explained 43.1% of the variance in the genetic data, and PC2 explained 23.2%. A Procrustes analysis comparing the first two principal components of the genetic data to the latitude-longitude coordinates of the sampling location indicated that there is significant geographic signal in the genetic data (Procrustes rotation 0.85, *p* < 0.001, **Fig. 4D-F**).

### Genetic analysis

Our genetic analysis revealed a pairwise F_ST_ value of 0.64, with an AMOVA showing that 63% of the variance was between species and 37% of the variance was within species. This agrees with the PCA analysis, where we see a separate cluster for the Eastern towhees (**Fig. 3E**).

## DISCUSSION

Since Oscine songbirds learn their songs from conspecifics, they are an excellent model system for studying how culturally transmitted traits affect evolution and speciation. Here, we examine variation in the learned songs of the Spotted and Eastern towhees, a sister species pair that each have broad ranges that span large distances, together covering ∼10 million km^2^. Since we have access to song recordings sampled throughout North America, we can detect variation that exists at these continental scales, allowing us to assess how traits have changed with distance. This pair of sister species have a relatively narrow zone of geographic overlap, where it has been noted that individuals occasionally hybridize (**Fig. S2**). Although hybridization occurs between these sister species—using eBird data, we estimate hybrids to comprise 0.4% to 1.6% of sightings in the zone of range overlap—our analysis suggested levels of genetic differentiation similar to those of other avian species (Irwin et al. 2018). This suggests that the two species are genetically distinct, indicating reproductive isolation in these sister species. However, we had few genotyped samples, and the majority of them were from the outer edges of the ranges, which limits our ability to assess genetic differentiation in the overlap zone. Nonetheless, these populations are considered separate species due to genetic, morphological, and song differences and seem to maintain themselves as separate species. Therefore, individuals are likely using specific traits or a combination of traits to recognize conspecifics, particularly because variation in song features exists not only between sister species but also within species. In this study, we analyze potential song features that differ between the Spotted and Eastern towhees, and we investigate whether song features could potentially allow individuals to reliably distinguish species-specific songs.

Our statistical analysis showed significant variation in song features across the longitude and latitude gradient overall, but showed stronger correlations and greater differences across longitude than latitude. For example, the number of syllables and bout duration of songs showed a decrease at around 120°W (**Fig. 3 A & B**). Additionally, the populations of Spotted towhees in the westernmost edge of the geographic range appear to have larger variation in some of the song features, including number of syllables and average syllable duration (**Fig. 3 B & C**); this pattern appears to correspond to the region where Spotted towhees have more rapid trills that occasionally were performed as a single syllable (with no separation between the repeated elements). Together, our results suggest that Spotted towhee songs differ predominantly in the westernmost part of their range, and geographic variation alone does not account for the song variation between Spotted and Eastern towhees. Overall, we found that no single song feature can be used to reliably distinguish between species’ songs. In contrast to most song features, the average syllable upper frequency seems to show a shift corresponding to the region where the two species’ geographic ranges meet (**Fig. 3 E & J**), but the range of frequencies overlaps between the Spotted and Eastern towhees. This song feature is also the highest loading in PC2, which captures a large portion of the variation in the song data. The highest loading for PC1 was average syllable duration, which was associated with the song differences observed in the westernmost Spotted towhees. The PCA revealed minimal separation in clustering using PC1 and PC2 (**Fig. 4B**). However, the song features of the two species were biased toward different portions of this cluster in PC-space, such that we could divide the plot into two sections and discriminate species with 73.6% accuracy, and a Procrustes analysis showed weak but significant geographic structure in the distribution of song data (**Fig. 4C**). After visualizing the song features with a UMAP projection, we found one primary cluster containing most of the Eastern and Spotted towhee songs, but, as in the PCA, each species primarily occupied a different portion of the cluster, and we could similarly partition the projection into two sections that could predict the species of each recording with 85.3% accuracy. Performing with even greater accuracy, our random forest classifier was able to correctly predict the species of 89.5% of songs, suggesting that several song features, in combination with one another or with morphology, could allow individual birds to reliably distinguish members of their own species. However, our random forest models trained on samples from the edges of the range were able to predict the species identification of songs from outside the zone of overlap better than those inside the zone of overlap (**Fig. 6B; Fig. S4**), supporting the notion that intermediate songs seem to exist (Sibley and West 1959). Our observation that songs of the Spotted and Eastern towhee are less distinguishable in the zone of overlap than at the edges of the ranges does not support the hypothesis that this sister-species pair has not developed increased song differentiation in this zone of overlap to discourage hybridization. Nonetheless, the incorrect predictions are distributed relatively evenly across the geographic range and not concentrated in the zone of overlap. In addition, the “hybrid/unsure” recordings—songs of potential hybrids and of towhees that were difficult for the recordist to categorize as Eastern or Spotted—did not fit a predictable longitude-based pattern of categorization by the random forest classifier.

Altogether, our analyses suggest that reinforcement of species boundaries is not readily detectable in towhee song, and other factors, such as cultural drift or differing habitats (e.g. vegetation density, altitude, climate), could also help explain the extensive amount of variation that we see in song features within and between species. For example, the acoustic adaptation hypothesis suggests that birdsong evolves under the constraints of the sound transmission properties of a given environment, and the songs are structured to increase the fidelity of song transmission in the native environment (Boughman 2002; Derryberry 2009; Morton 1975). As such, individuals with certain song traits may be more fit in specific environments if their song is able to transmit through the environment and be more easily detected. Additionally, as individuals transcend the zone of overlap into the other species’ range, female preference for certain traits could lead to differential fitness in males depending on their songs (Price 2007). For example, if females prefer mates with traits that fall towards the extremes, those with intermediate songs would have decreased fitness. This may contribute to constraining the width of the hybrid zone and maintaining reproductive isolation. As such, future studies incorporating environmental characteristics at the population level would give us greater insight into factors affecting the evolution and speciation of avian species. Further, analyses that account for geographic distance between songs could help assess whether songs accumulate changes by cultural drift, fitting an isolation-by-distance pattern. Additionally, differences in the trills of the Spotted and Eastern towhee may be features that are important for species recognition. A study by Richards (1981) suggests that the trills of the northeast populations of Eastern towhees function as a messaging component and may indicate species recognition. Therefore, future analyses could compare only the trills of Spotted and Eastern towhee songs to determine whether these trills have more species-specific differences than the songs as a whole. Finally, using playback experiments in the field to assess how individuals respond to songs of conspecifics versus songs of heterospecifics would directly test whether individuals can discriminate between songs and whether a preference exists for certain song features.

In sum, we find that subtle song differences between Eastern and Spotted towhees can lead to relatively reliable species distinctions that are detectable with a machine-learning classifier trained on multiple song features but not with simple statistical comparison of single song features. However, it is interesting to note that these distinctions are less apparent in the zone of species overlap, where they have been hypothesized to be most useful. Based on these results, we hypothesize that there is not a strong selection pressure in the zone of overlap favoring song differentiation to limit hybridization below its current level of ∼1%. Perhaps since this zone of overlap is relatively sparsely populated with both Spotted and Eastern towhees, the limited opportunity for potential breeding interactions between species does not lead to a strong advantage for male birds that have reliably distinguishable songs or for the female birds that prefer them.

## Author Contributions

X.L.D. and N.C. conceived and designed the project. X.L.D. downloaded and parsed song recordings to produce song-feature data. X.L.D. wrote analysis code, ran all analyses, and created the figures and tables with assistance from N.C. X.L.D. and N.C. wrote the manuscript.

## Data and materials availability

All analysis code and data are available at https://github.com/CreanzaLab/TowheeAnalysis, including metadata about each song recording.

