## Supplemental Tables and Figures for "The role of learned song in the evolution and speciation of Eastern and Spotted towhees"

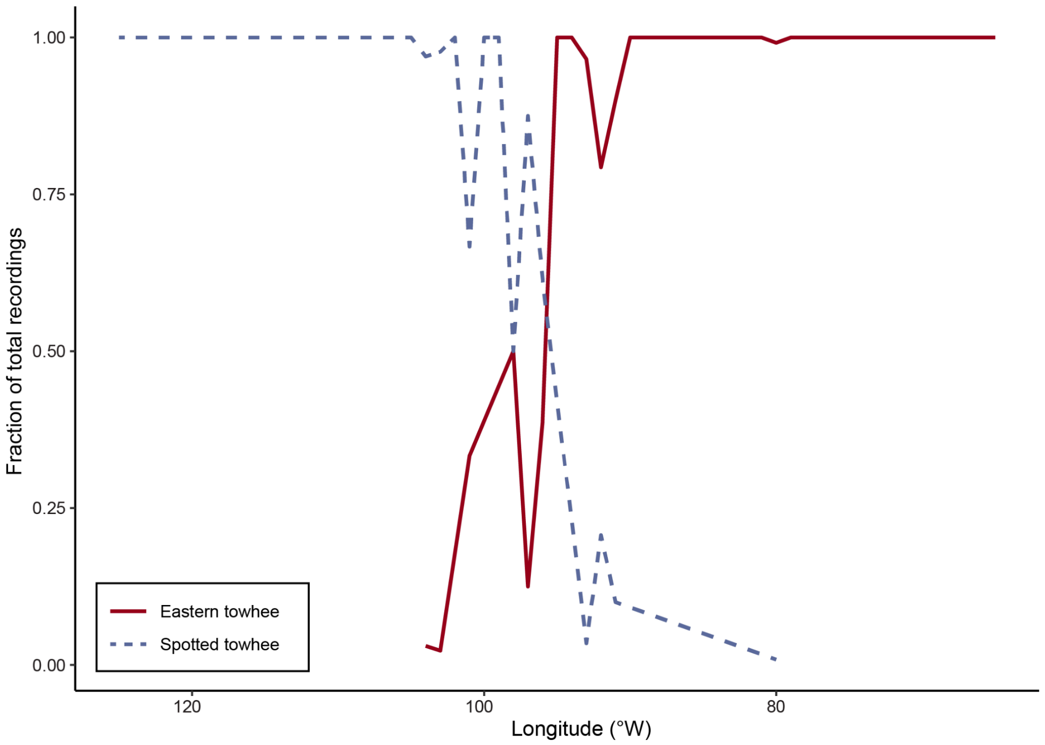


**Supplementary Figure S1.** Line plot of the fraction of Spotted towhee and Eastern towhee recordings, calculated as the number of recordings of each species divided by the total number of recordings of either species across North America during the breeding season ($N_{total}$=3037; $N_{Spotted\_towhee}$=1146; $N_{Eastern\_towhee}$=1891). This plot was used to determine the zone of overlap (102°W - 91°W).

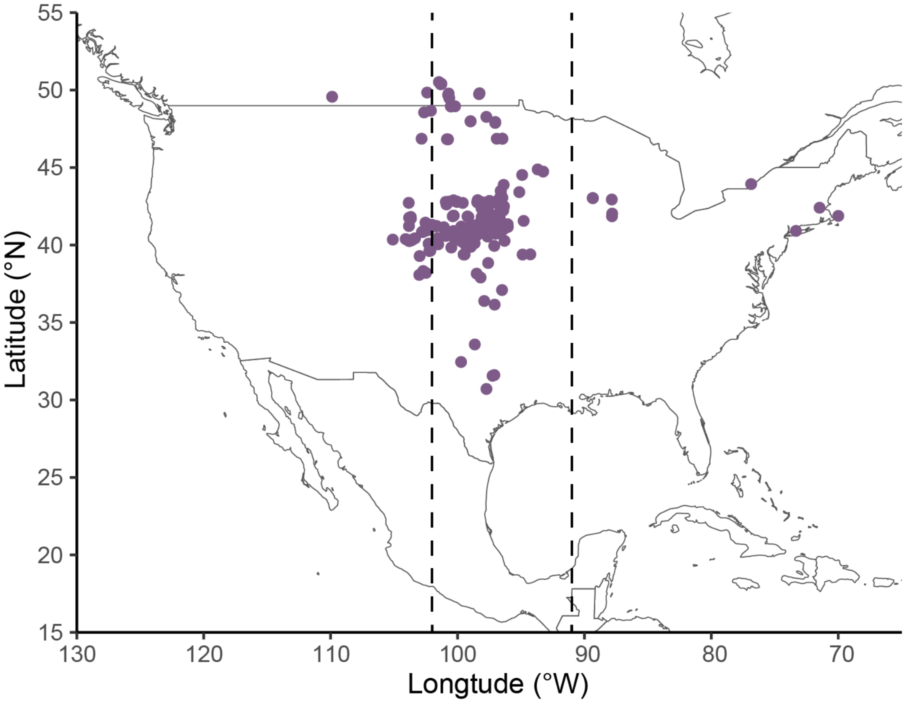


**Supplementary Figure S2.** Map of putative Spotted towhee x Eastern towhee hybrid sightings (N = 203) during the breeding season. The dotted line represents the zone of overlap determined by the co-occurence of song recordings (102°W - 91°W). Data obtained from eBird.

| Song Feature | Spearman | | | | | | Wilcoxon | |
|---|---|---|---|---|---|---|---|---|
| | Latitude | | | Longitude | | | | |
| | S | *p*-value | Spearman's ρ | S | *p*-value | Spearman's ρ | W | *p*-value |
| bout duration (ms) | 4629660239 | 0.646 | 0.008 | 2494939142 | 2.94E-163 | 0.466 | 465055 | 1.19E-153 |
| number of syllables | 4989982768 | 1.46E-04 | -0.069 | 4858672750 | 0.025 | -0.041 | 1237365 | 4.66E-11 |
| number of syllables per bout duration (1/ms) | 4931072202 | 1.94E-03 | -0.056 | 5629074782 | 2.19E-30 | -0.206 | 1460543.5 | 2.75E-58 |
| largest syllable duration (ms) | 4356526342 | 2.28E-04 | 0.067 | 3471956918 | 9.25E-47 | 0.256 | 596573.5 | 5.30E-96 |
| smallest syllable duration (ms) | 4370373624 | 4.28E-04 | 0.064 | 4647120062 | 0.800 | 0.005 | 1015487 | 3.67E-03 |
| average syllable duration (ms) | 4317377652 | 3.33E-05 | 0.075 | 3939282376 | 4.79E-18 | 0.156 | 765409 | 5.11E-42 |
| number of unique syllables | 4442783805 | 7.69E-03 | 0.048 | 2313217519 | 7.39E-196 | 0.505 | 390954.5 | 8.65E-205 |
| number of syllables per number of unique syllables | 5059061416 | 3.92E-06 | -0.084 | 5940866796 | 7.30E-53 | -0.273 | 1574420.5 | 1.06E-97 |
| average syllable upper frequency (Hz) | 5267647290 | 1.26E-12 | -0.128 | 6428804873 | 3.37E-103 | -0.377 | 1753604.5 | 5.49E-180 |
| average syllable lower frequency (Hz) | 4685328208 | 0.843 | -0.004 | 5411845292 | 1.08E-18 | -0.159 | 1371763 | 8.51E-35 |
| maximum syllable frequency (Hz) | 5310111500 | 2.83E-14 | -0.137 | 5081212132 | 1.07E-06 | -0.088 | 1355688.5 | 3.28E-31 |
| minimum syllable frequency (Hz) | 4744796056 | 0.368 | -0.016 | 6491094077 | 3.97E-111 | -0.390 | 1617674.5 | 3.42E-115 |
| overall syllable frequency range (Hz) | 5251685729 | 4.93E-12 | -0.125 | 4269177314 | 2.35E-06 | 0.086 | 1109977.5 | 0.259 |
| largest syllable frequency range (Hz) | 5207820726 | 1.72E-10 | -0.116 | 5299230684 | 7.67E-14 | -0.135 | 1374444.5 | 2.02E-35 |
| smallest syllable frequency range (Hz) | 5089768974 | 6.35E-07 | -0.090 | 6536247306 | 4.14E-117 | -0.400 | 1729314 | 1.96E-167 |
| average syllable frequency range (Hz) | 5153333131 | 9.72E-09 | -0.104 | 5915472064 | 9.08E-51 | -0.267 | 1558821.5 | 1.54E-91 |

**Supplementary Table S1.** Spearman's rank correlation ($\alpha_{adjusted}$=0.00156) and Wilcoxon rank sum test ($\alpha_{adjusted}$=0.003125) of Spotted towhee and Eastern towhee song features ($N_{Spotted\_towhee}$=1146; $N_{Eastern\_towhee}$=1891) using the subset of samples recorded during the breeding season.

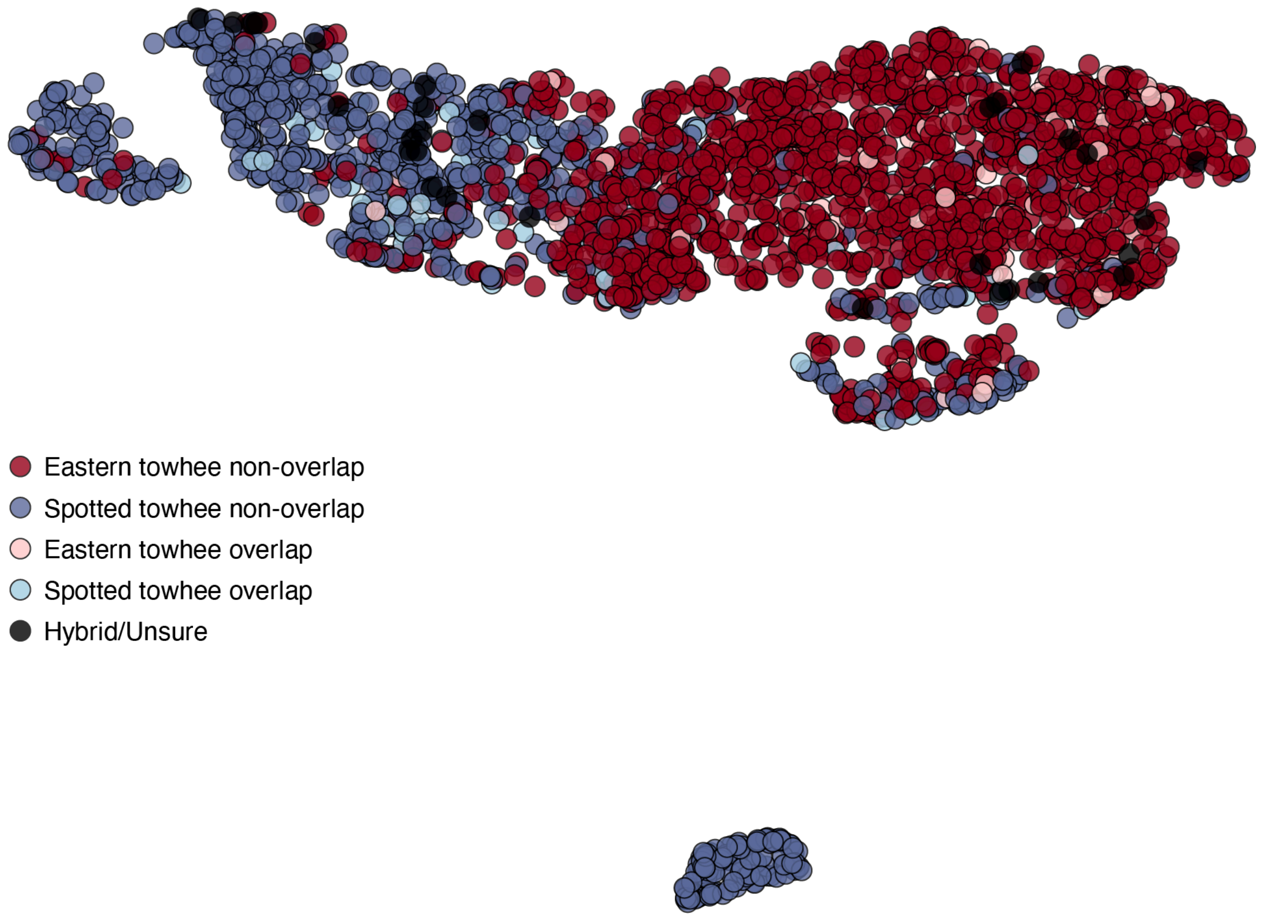


**Supplementary Figure S3.** UMAP projection of Eastern and Spotted towhee song-feature data using the subset of samples recorded during the breeding season. Each point represents an analyzed song bout ($N_{total\_bouts}$=3037; $N_{Spotted\_towhee}$=1146; $N_{Eastern\_towhee}$=1891), with Eastern towhee songs shown in shades of red and Spotted towhee songs in shades of blue. The lighter colors represent recordings from the zone of species overlap. Black dots indicate the 30 recordings from individuals that were classified as potential hybrids ("hybrid/unsure"). Using a linear discriminant classifier to partition the UMAP projection into two sections, we could accurately predict the species of 85.7% of recordings (n_neighbors=15 and min_dist=0.1). Using values of n_neighbors up to 50 and values of min_dist up to 0.9, the linear classifier showed prediction accuracies ranging from 79.6% to 85.7%.

| Model | Training Data | Number of Trees | Prediction Accuracy |
|---|---|---|---|
| A | All bout samples | 100 | 89.2% |
| B | All bout samples | 500 | 89.5% |
| C | All bout samples | 1000 | 89.4% |
| D | Zone of non-overlap | 100 | 88.0% (non-overlap)<br>81.3% (overlap) |
| E | Zone of non-overlap | 500 | 89.3% (non-overlap)<br>82.6% (overlap) |
| F | Zone of non-overlap | 1000 | 88.3% (non-overlap)<br>81.6% (overlap) |

**Supplementary Table S2.** Prediction accuracies of random forest models trained on 16 song features from samples of Spotted towhees and Eastern towhees using different numbers of decision trees. (A-C) Predictions of subset of all song samples ($N_{test}$=879) trained on song data from the entire geographic range ($N_{Spotted\_towhee}$=1062; $N_{Eastern\_towhee}$=1062). (D-F) Models trained on a subset of samples obtained from the non-overlap zone ($N_{Spotted\_towhee}$=1062; $N_{Eastern\_towhee}$=1062). The model was tested on a random subsample of song bouts from both the zone of non-overlap ($N_{test\_nonoverlap}$=299) and the zone of overlap ($N_{test\_overlap}$=299). Increasing the number of trees did not improve the accuracy of the models' predictions. We report the results from models with 500 trees in the main text.

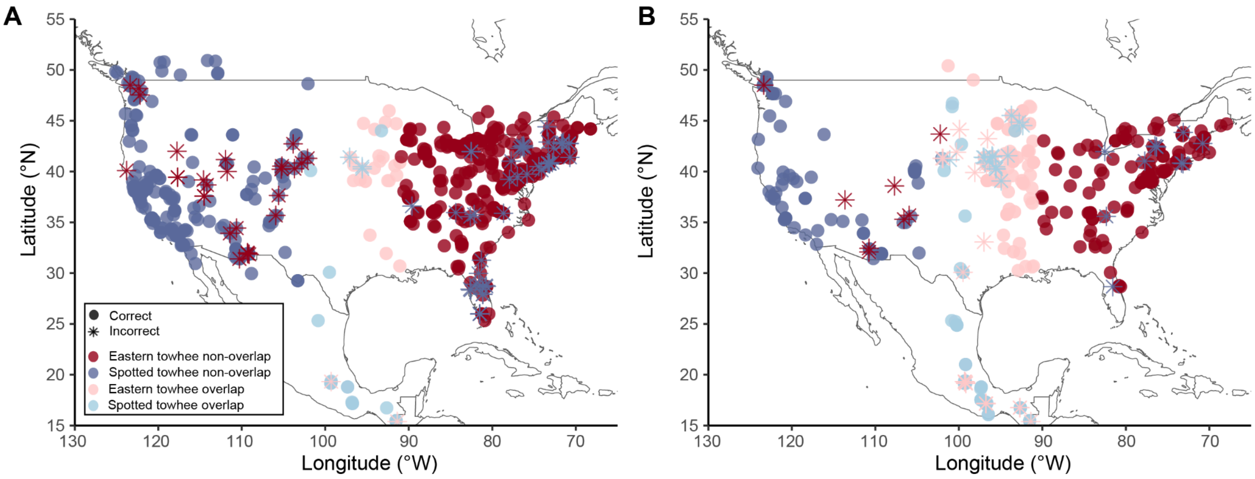


**Supplementary Figure S4.** Geographic distribution of species predictions using a random forest model trained on 16 song features from only those samples of Spotted towhees and Eastern towhees that were recorded during the breeding season. (A) We trained a model on song data from the entire geographic range of songs recorded during the breeding season ($N_{Spotted\_towhee}$=853; $N_{Eastern\_towhee}$=853) and tested how well it predicted the species identification of a subset of breeding-season song samples ($N_{test}$=760; accuracy = 88.9%). (B) We then trained a second model on a subset of samples obtained during the breeding season from the non-overlap zone ($N_{Spotted\_towhee}$=853; $N_{Eastern\_towhee}$=853). This model was tested on a random subsample of breeding-season song bouts from both the zone of non-overlap ($N=_{test\_nonoverlap}$=211; accuracy = 91.0%) and the zone of overlap ($N_{test\_overlap}$=211; accuracy = 82.9%).

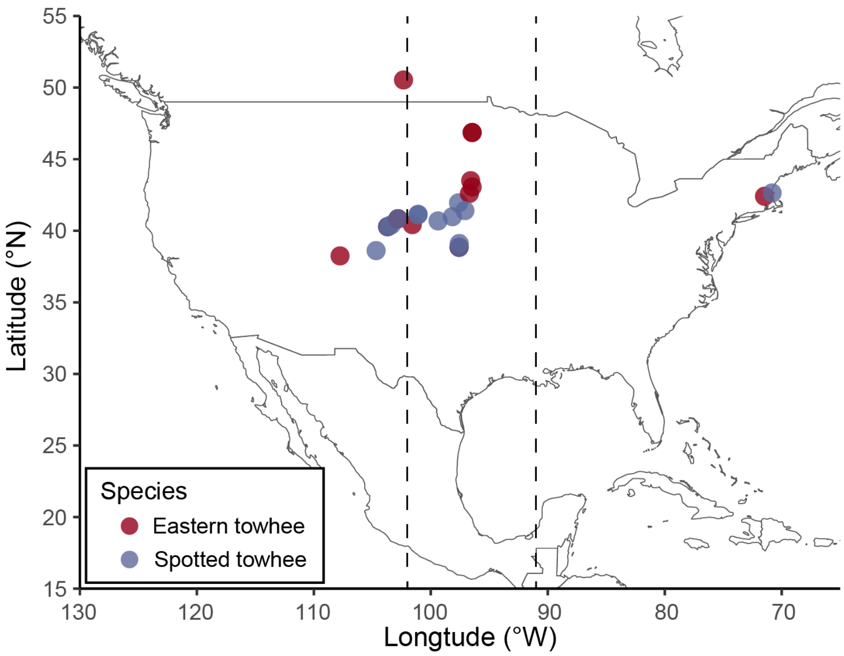


**Supplementary Figure S5.** Geographic distribution of random forest model predictions of species identity of song bouts from recordings of "hybrid/unsure" towhees ($N_{predict}$=31). The model was trained on 16 song features from a subset of samples of Spotted towhees and Eastern towhees obtained only from the non-overlap zone during the breeding season ($N_{Spotted\_towhee}$=853; $N_{Eastern\_towhee}$=853). The model predicted that 19 of these "hybrid/unsure" recordings were Spotted towhees and 12 were Eastern towhees, with no discernable longitudinal gradient in the predictions. The dotted line represents the zone of overlap determined by the co-occurence of Eastern towhee and Spotted towhee song recordings (102°W - 91°W).
